# Drug prediction for reversing AD/PD transcriptional profiles using an aging of systems-centric approach

**DOI:** 10.1101/2022.11.01.514657

**Authors:** Gabriela Bunu, Dmitri Toren, Eugen Ursu, Simona Ghenea, Vadim E. Fraifeld, Robi Tacutu

**Affiliations:** Systems Biology of Aging Group, Institute of Biochemistry of the Romanian Academy, Bucharest, Romania; The Shraga Segal Department of Microbiology, Immunology and Genetics, Center for Multidisciplinary Research on Aging, Ben-Gurion University of the Negev, Beer-Sheva, Israel; CellFabrik SRL, Bucharest, Romania

**Keywords:** aging, Alzheimer’s disease, Parkinson’s disease, bioinformatics

## Abstract

Age-related pathologies are so widely presented in old age that in most cases they are hardly distinguishable at the molecular level from the so-called ‘‘normal’’ aging. Both aging and age-related diseases are characterized by a wide range of transcriptional and epigenetic changes that underlie the physiological or pathological phenotype, with plenty of overlap in their signatures, but also with differences. In most pathological conditions it is rather the dysregulation of a complex network of genes than a problem with a single gene dysregulation that causes its emergence or progression, and aging differently gives a “predisposition” towards an age-related pathology or another, or in a favorable situation towards none. The important question is how similar are the transcriptional changes during “healthy” aging with those that occur in age-related diseases. In this study, we explore gene expression data to answer this question and aim to predict which drugs and compounds could have a reversing effect on their common drift.

## Introduction

Despite pathologies being intimately connected to aging itself, the exact molecular relationship between aging and age-related diseases (ARDs) still remains an unsettled issue in biogerontology (Budovsky et al., 2006; Hekimi, 2006; Yang et al., 2016). The frequency of major life-threatening degenerative pathologies in humans, including atherosclerosis, cancer, neurodegeneration, type 2 diabetes, osteoporosis, and sarcopenia, progressively increases in the post-reproductive period (Cutler and Mattson, 2006). Some pathways are common between aging and ARDs (Budovsky et al., 2009; Wolfson et al., 2009; Tacutu et al., 2011; Fernandes et al., 2016), however this still does not explain why some people appear to be suffering from one or more age-related pathologies from a younger age while others age in a “healthy” manner, showing no clear indication of any ARDs until their death. Thus far, hundreds of genes have been identified as being involved in aging, longevity, and ARDs (Tacutu et al., 2010, 2013, 2018), and many common genes to both aging/longevity and ARDs have been shown to act in a cooperative manner, forming entire gene networks (Budovsky et al., 2007; Tacutu et al., 2010a, 2010b; Wolfson et al., 2009). This led to the idea that ARDs are so widely presented in old age that they are hardly distinguishable from the so-called ‘‘normal’’ aging and, in fact, represent its diverse manifestations, being an essential component of the aging process (Budovsky et al., 2007).

While it is highly probable that some common molecular mechanisms stand behind both aging/longevity and ARDs, some evidence has shown that the aging processes could also diverge at different points, leading the healthy aging system towards the development of one pathology or another (Demetrius et al., 2014; Driver, 2014).

Overall, aging and ARDs are characterized by a wide range of transcriptional and epigenetic changes that underlie the physiological or pathological phenotype. Recent studies show that in most pathological conditions it is rather the dysregulation of a complex network of genes than a problem with a single gene dysregulation that causes its emergence or progression (Smith and Flodman, 2018). So, the presence of common genes and networks is essential but still not sufficient evidence for a common molecular basis for aging and ARDs. The important question is whether there is similarity between the transcriptional drift during “healthy” aging and that which occurs across ARDs. Perhaps even more importantly, another question is whether we can derive signatures of transcriptional changes that could still be reversible. In this paper we focus on a multi-study analysis of aging and ARD microarray datasets that could explore the above hypothesis.

## Results

### Aging and age-related disease datasets

The first step in this study was the collection of human microarray datasets relevant to healthy and pathological aging. For healthy aging, transcriptional datasets were considered if the authors reported a comparison between younger and older individuals, with both compared groups being considered healthy (or more accurately, without showing any signs of age-related pathologies). Datasets of interest were considered those that contained samples from different age segments, grouped by us mainly into three categories - young (15-35 years old), middle-aged (35-65 years old), and old (65-89 years old). For the purpose of the current study, dataset samples from nonagenarians, centenarians and supercentenarians were excluded. While generally there is extraordinary value in such datasets, we believed the transcriptional profiles of long-lived humans, at advanced ages, would provide mixed signals into our analysis. On one hand, these samples reveal the transcriptome at a very advanced age, which should entail stronger signals about the age-related differences, but on the other hand, they also contain the beneficial transcriptional differences that allowed long-lived individuals to reach such advanced age.

For age-related diseases, the following pathologies were included in the search: Alzheimer’s Disease (AD), Parkinson’s Disease (PD), Type 2 Diabetes, Atherosclerosis and Osteoporosis. In total, 54 microarrays were selected, 24 datasets for association with healthy aging and 30 datasets for pathologies: 6 for AD, 10 for PD, 2 for Type 2 Diabetes, 4 for Atherosclerosis and 8 for Osteoporosis. For the selected datasets, sample conditions were defined and a list of comparisons of interest was compiled. The full list of 54 datasets can be found in Supplementary Table 1.

### Similarity between transcriptional changes in aging and age-related diseases

For all selected datasets, samples were normalized and for the comparisons of interest, differential gene expression was computed resulting in a series of transcriptional profile shifts (differential expression changes between two experimental conditions). The full list, briefly describing the 104 transcriptional shifts/comparisons can be found in Supplementary Table 2. In order to evaluate which of these shifts are changing the transcriptome in the same direction and which are opposite, we next computed a similarity score between any two shifts, using an *in-house* developed Python script (which is further detailed in the Materials and Methods section). The script was then used to compute all pairwise similarities and to construct an undirected network (included in Fig. 1A) with the score representing the strength of an interaction between nodes (the transcriptional profile shifts).

**Figure 1.**
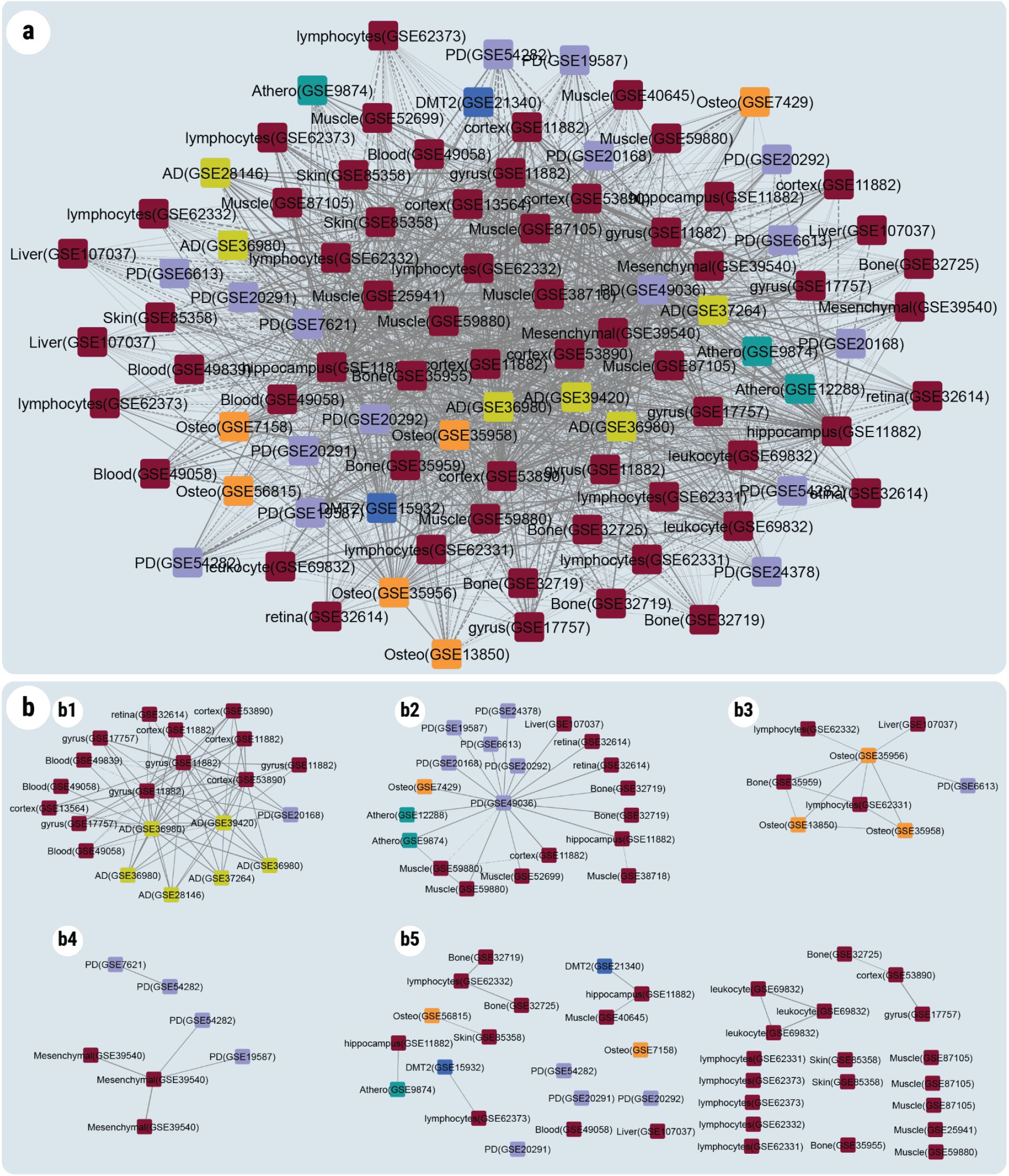
**A**. Undirected network of similarities between any two transcriptional shifts, selected from the studied datasets. **B**. Clusters obtained using the MCL algorithm, applied to the network shown in panel A, and using the similarity scores as strength for the edges. **A-B**. The network and clusters include the following comparisons: 1) “healthy” aging (old vs young) - maroon; 2) AD - yellow; 3) PD - purple; 4) atherosclerosis - cyan; 5) type 2 Diabetes - dark blue; 6) osteoporosis - orange. For schematic simplification, the displayed titles include only the tissue and the name of the GSE - each node represents in fact one comparison (transcriptional shift) between two analyzed conditions in that dataset/study. Edges between nodes represent similarity scores between two comparisons. Continuous line - positive scores (similar shifts); dashed line - negative scores (opposite shifts).

### Clustering of system modifications in physiological and pathological aging space

Using a clustering algorithm, the network in Fig. 1A was transformed to identify groups of pathophysiological causes that lead to relatively similar change patterns (Fig. 1B). In total, 4 clusters were obtained (b1-b4), with a series of additional 2-3 node groups (b5 area) which probably characterize aspects more private to certain conditions, and less common to all forms. As can be seen in panel B, the two main clusters in the top-left of the figure were better outlined. First cluster (denoted in the figure as “b1”) includes neurodegenerative changes - mainly characteristic to AD (yellow nodes), and to several aging-related comparisons - especially in brain regions, but also a few in blood. In total, the first cluster (referred in this article as Cluster-AD) has 21 nodes (transcriptional comparisons), including 14 old vs young comparisons (from 7 studies), and 6 AD vs control (from 4 studies). In general, the similarity scores show consistency between datasets (121 positive, and only 8 negative scores), suggesting a strong consensus. Interestingly, Cluster-AD also includes a transcriptional shift reported in one of the studies of Parkinson’s Disease.

The second cluster (denoted in the figure as “b2”) is more characteristic to changes that occur in Parkinson’s Disease (the purple nodes), with a peculiar similarity to several pattern changes that appear in Atherosclerosis and Osteoporosis. For this cluster (referred in this article as Cluster-PD), a wider distribution of tissues is observed for aging-related changes - including brain, but also muscle, bone, retina and liver. It contains 20 nodes and 23 edges. The nodes belong to 11 old vs young comparisons (7 different studies), 6 PD vs control (6 studies), 2 atherosclerosis vs control (2 studies) and 1 osteoporosis vs control. Only 2 similarity scores are negative. Cluster-PD has a lower density than Cluster-AD and there is a central hub node that is highly similar (score > 0.6) to all but one other node.

The next obvious question is what is the consensus in the transcriptional shift for each of the two clusters, i.e. what common and consistent (or almost common/consistent) changes are the core drivers of these age-related processes - either seen in pathological form after a significant advance or perhaps in a pseudo-physiological form as predisposing states of the aging system. To answer this, we next performed a meta-analysis for each cluster, considering all the different datasets in a cluster. In order to have a more inclusive analysis, in which small-size effects are still considered and in which the focus is more on the consistency of direction rather than the strength of the signal (which in some studies might be limited by the available sample size) we have performed a custom-designed meta-analysis. This analysis took into consideration all gene expression changes at p-value < 0.05, without taking into account the FDR multiple testing correction, and ensured that the genes selected as up- or down-regulated show a fold change > 2 in at least one dataset, while no other dataset shows a significant change in the opposite direction (the steps of the meta-analysis are described in Materials and Methods).

For comparison, we have also carried out another meta-analysis, using the RankSum analysis from the RankProd library, with FDR-adjusted data from each dataset (FDR <0.05, FC increase/decrease >= 50%; data not included). It should be mentioned that while the results from our custom analysis are more lax (resulting in larger lists), the functional enrichment at the level of biological processes and signaling pathways showed the results being relatively similar. The main difference between the two approaches is that for the RankSum with FDR, the number of contributing datasets to the analysis is drastically reduced because most of the comparisons from public datasets have limited statistical power, and weak signals cannot be clearly distinguished even for a 50% FC threshold. For example, for Cluster-AD, only 8 out of 21 comparisons show any significant differences by themselves - thus reducing the input for the meta-analysis to only 4 aging datasets and 2 AD datasets (instead of 7 and 6, respectively). By contrast, our meta-analysis criteria led to all 21 transcriptional comparisons being included in the selection. As a result of the meta-analysis, the list of genes consistently changed across Cluster-AD was reduced to 410 up-regulated genes and 833 down-regulated genes. For Cluster-PD, the list included 318 up- and 229 down-regulated genes. Supplementary Table 3, contains these lists for both clusters.

### Functional characterization of the main two clusters

To further understand the characteristics of up- and down-regulated genes from each cluster, we conducted a functional module analysis, using the HumanBase online tool (Greene et al., 2015). The analysis revealed that in the Cluster-AD (Fig. 2a), upregulated genes form five functional modules that are involved in viral response and interferon-gamma (MA1), cellular response to zinc ion (MA2), bacterial immune response (MA3), and morphogenesis (MA4), response to peptide and microtubule cytoskeleton organization (MA5). In contrast, downregulated genes from Cluster-AD (Fig. 2b), generated seven functional modules that are related to dendrite regulation (MB1), neuron projection and differentiation (MB2), aspartate metabolic processes and actin rod assembly (MB3), potassium ion transport (MB4), morphogenesis (MB5), and anion transport (MB6).

**Figure 2.**
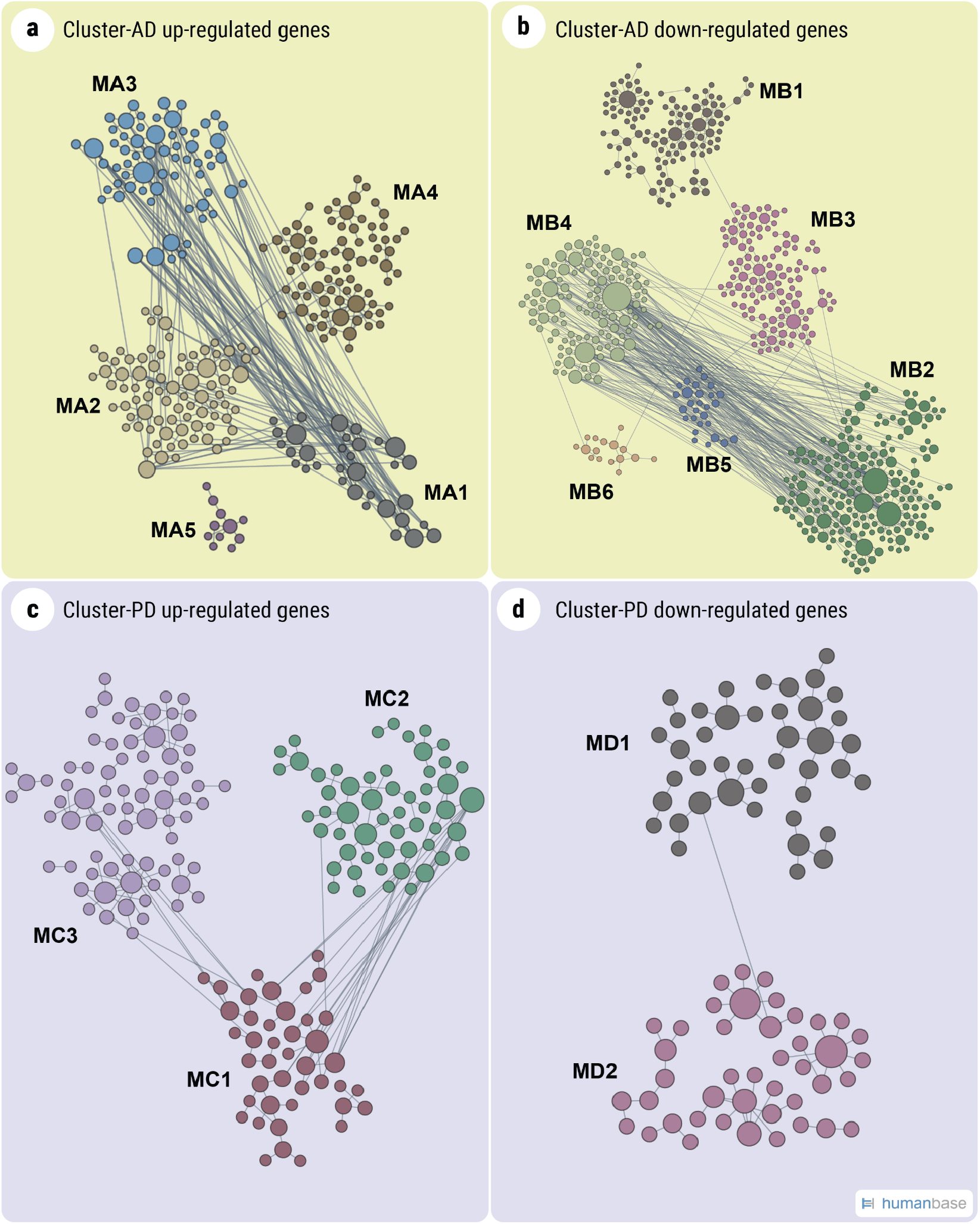
Functional module network of the up-and down-regulated genes from the cluster analysis. The network was built for the lung tissue, using HumanBase. The interaction network is built using the closest gene neighbors and then clustered based on enrichment in GO categories. **A**. Cluster-AD up-regulated genes five modules. **B**. Cluster-AD up-regulated genes six modules. **C**. Cluster-PD up-regulated genes three modules. **D**. Cluster-PD up-regulated genes two modules.

The 3 modules based on Cluster-PD’s upregulated genes (Fig. 2c) are primarily reflecting RNA polymerase activities (MC1), protein monoubiquitination (MC2), and response to cAMP (MC3). Downregulated genes from Cluster-PD on the other hand encompass only 2 modules (Fig. 2d). Module MD1 is involved in the regulation of microtubule polymerization and apoptotic signaling pathway. Similar to the MB2 module in Cluster-AD, module MD2 is enriched in the regulation of neurogenesis, neuron differentiation and other neuron-related processes.

Next, we also performed a gene set enrichment analysis (GSEA – prerank) on the ranked list of genes from the meta-analysis in order to further characterize the function space of the genes comprising Cluster-AD and Cluster-PD. The full list of results is available as Supplementary Table 4. Enrichment was examined with respect to several relevant databases: GO Biological Processes, KEGG, Reactome and ChEA. For Cluster-AD, we observed significant enrichment suggestive of upregulation of general immune system terms (innate immune system, neutrophil degranulation, interferon gamma response), as well as some more specific immune system terms (FOXO1, NCOR, IL-8, IL-6/JAK/STAT3, IL-2/STAT5). Terms related to infections (Epstein-Barr virus infection, Coronavirus disease) were also observed. Terms not related to the immune system which were observed to be enriched included: hypoxia, regulation of angiogenesis, phagosome and apoptosis. With regards to downregulation, we consistently observed terms related to chemical synapses: transmission across chemical synapses, GABAergic synapses, glutamatergic synapses and cholinergic synapses. 6 significantly downregulated gene sets were related to the SUZ12 protein.

Surprisingly, we did not identify any significant enrichments based on the results from Cluster-PD after FDR adjustment.

### Reversing the transcriptional drifts associated with neurodegenerative aging

Considering the sources of expression changes that contribute as the input for the meta analysis, namely aging and neurodegenerative disease shifts, it is tempting to suggest that the common changes (i.e., the meta-analysis results) might also have a determinant role in the degenerative progression of an aging system, either pushing it to a pathological state (such as AD or PD), or making it more vulnerable even though seemingly healthy (as in “healthy” aging). Idealistically, a therapeutic approach to tackle this would be to target and reverse the entire transcriptional drift, however, this seems highly improbable to achieve. Instead, a search for drugs and compounds that move the system state as much as possible towards the initial state might be attainable.

ConnectivityMap is a tool that allows users to search through a large database of perturbational studies (based on *in vitro* data), identifying the genetic and pharmacologic perturbagens that cause similar or, on contrary, opposite signatures. As such, we next used the topmost up- and down-regulated signatures (sorted by their fold change) from the meta-analyses done on clusters 1 and 2, to identify the compounds with the most negative scores.

From the analysis of Cluster-AD 15,076 significant (FDR<0.05) potential compounds resulting in an opposite effect to the transcriptomic profile determined by aging/neurodegenerative pathologies were identified. Among these compounds, we found some drugs whose mechanisms of action were previously associated with aging and aging-related processes. For example, manually going through the compounds with the 200 top-most negative scores, we identified 9 drugs (out of a full list of 136,460 compounds in the ConnectivityMap data bank and 26,557 drugs with FDR<0.05) that could reverse the consensus changes induced by the conditions of Cluster-AD and had been already associated either with aging or with some aspects of the aging process.

By contrast, for Cluster-PD we have identified only a list of 194 significant drugs (FDR<0.05) with negative similarity scores. Nevertheless, among these drugs we also found 8 drugs that could reverse its consensus transcriptional change and which were previously associated with aging-associated processes or age-related pathologies.

Since the drugs in Cluster-AD/Cluster-PD are characterizing an entire cluster and are not specific only to a certain pathology, we hypothesized that some of them might work in a broader way, via mechanisms of reprogramming and induced pluripotency. Recently we compiled a manually curated list of 92 chemical compounds that can either induce or enhance pluripotency, alone or in combination with TFs (Knyazer et al., 2021). Indeed, many of these small molecules can be found with negative scores in our current profile-reversing predictions. For example, 21 of the 92 compounds were found to have negative scores in the analysis of Cluster-AD, and 18 of them had significant values (FDR<0.05). Some of these potential targets are discussed in the next section. On the other hand, the drugs that could potentially reverse the signature of Cluster-PD did not include any of the re-programming drugs, pointing out that a brain rejuvenation solution would work differently for PD than for AD.

## Discussion

Up to date, comparisons of aging signatures between different tissues have been carried out (e.g. Zahn et al. 2006), although in most cases limited to 2 or 3 tissues. The authors came to the conclusion that there is a common aging signature between skeletal muscle, brain and kidney, that comprises several pathways: ECM genes, cell growth genes, and complement activation genes which significantly increase their expression with age, and the chloride transport genes and electron transport genes, which significantly decrease their expression with age in all three human tissues (Zahn et al. 2006). However, the conclusion regarding a common aging signature was based on the comparison of pathways but not of the individual genes involved. Thus, the question as to the existence of a common aging gene expression signature remained open. In another previously reported study (Rodwell et al., 2004), age-related expression changes were compared between kidney and skeletal muscle (Welle et al., 2003), however the authors did not find any similarity directly in aging signatures between the two tissues. Later, in a meta-analysis of 27 datasets from mice, rats and humans, several genes were found to be consistently changed with age (56 up and 17 down), involving mostly inflammation, immune response, the lysosome, collagen genes, energy metabolism (particularly mitochondrial genes), and cellular senescence (de Magalhães et al., 2009). None of these studies however targeted directly the involvement of aging in ARDs. Moreover, while some research has been done to identify specific gene signatures for ARDs, much still remains to be done before gene expression could be used as a biomarker before the actual symptoms. For example, Smith and Flodman, pointed out in their review the lack of gene expression research despite the fact that numerous diseases have been linked to genetic or genomic defects (Smith and Flodman, 2018). Lastly, in terms of integrative aging/ARDs analyses, there is only a limited amount of systems biology studies and the molecular interactions between different processes are rarely evaluated in a systems biology manner, as we did for example in Tacutu et al., 2011. In this work, we looked comparatively at how the transcriptional profile shifts during aging and as a consequence of ARDs, and tried to understand what aspects of ARDs are similar to aging, and to each other. The results showed two potentially divergent drifts (or pathological “manifestations” of aging), characterized by the changes in the two separate clusters of our analysis - the Cluster-AD and Cluster-PD. It should be acknowledged that our analysis is dependent on the availability of data, and it is possible that, with the appearance of more datasets, the meaning of the clusters might become more general. For the time being however, it seems that the two main clusters are specific to AD and PD.

The HumanBase functional module analysis of the clusters (Fig. 2) showed, as expected, some enriched up- or down-regulated processes that are in accordance with the two pathologies. For example, AD is known to be linked to the immune system (Bettcher et al., 2021), one of the recurring processes in our analysis. The role of zinc in AD was also previously reported (Watt et al., 2011) in agreement with cellular response to zinc ion being upregulated in Cluster-AD. Dysregulated protein phosphorylation, microtubule cytoskeleton organization and morphogenesis are also determining conditions in brain aging and AD (Ferrer et al., 2021). In both diseases, AD and PD, neurogenesis and neuronal regulation, were down-regulated. RNA polymerase activity was reported to play a critical role in AD (Kang and Shin, 2015; Majidinia et al., 2016). Monoubiquitylation regulates processes that range from membrane transport to transcriptional regulation (Hicke 2001) and there have been shown that monoubiquitination might play an important role in Lewy body formation (Engelender 2008). cAMP, whose response was upregulated in Cluster-PD is a potent regulator of innate and adaptive immune cell functions and its signaling was previously linked to PD (Santini et al., 2008; Goto 2017).

Similarly, the GSEA enrichment results for Cluster-AD highlighted well-known associations of aging, especially with immune system processes, such as the well-characterized connection between aging and FOXO1 (Martins et al., 2016). Infection-related terms are commonly occurring in enrichment analysis on aging datasets, and here we see no exception. This is not surprising, as many of the genes involved in infection processes overlap with genes associated with immune processes and with age there is a known shift towards immunosenescence (Wang et al., 2022). Additionally, there is already some data associating the development of AD to infections, such as with EBV (Zhang et al., 2022). We also observed downregulation of many synapse-related terms, which might drive broader, widespread aging-related alterations. On the other hand, the absence of significant enrichments for Cluster-PD may suggest significant functional heterogeneity of the genes included in this cluster.

The most important part of the methodology that we used in this study is whether it could be used to actually predict new drugs or other compounds that might reverse the profile shift that occurs during aging and ARDs. Using the clusters’ signatures, we found several drugs for each cluster that were indeed previously associated either with aspects of aging or the disease.

### Drugs identified from the analysis of Cluster-AD

AMG-487 (CC chemokine receptor antagonist), BRD-K68144790 (apoptosis stimulator), and diethylcarbamazine (lipoxygenase inhibitor) were found to have opposite signatures to that of Cluster-AD. Other examples include selumetinib, a MEK inhibitor, for which it is known that MEK inhibition can be used in the treatment of some forms of cancer and in the prevention or suppression of cellular aging (Steelman et al., 2011).

Heat Shock Proteins (HSPs) are known to be pro-longevity and can improve proteotoxicity associated with aging and have a regulatory role in cellular senescence, apoptosis and cancer (Tower, 2009), and homosalate, an HSP inducer was also identified with a highly dissimilar transcriptional change compared to the signature of Cluster-AD.

GDC-0152, an inhibitor of XIAP (X-linked inhibitor of apoptosis), and AZD-8055, an inhibitor of the well-known longevity-associated gene mTOR were also identified among the drugs with lowest scores. PPAR proteins are important regulators in various pathophysiological processes associated with age - especially processes involved in energy metabolism and oxidative stress (Erol, 2007). PARP1 for example can be considered a molecule with a pleiotropic antagonistic effect - on the one hand it protects cells from senescence in physiological conditions, on the other hand PARP1 supports cell death and functional decline in aging and pathophysiological conditions (Mao and Zhang, 2022). Pioglitazone, an agonist of PPAR receptors, and rucaparib - another PARP inhibitor also scored negative.

### Drugs identified from the analysis of Cluster-PD

Several drugs already approved for diseases included in Cluster-PD or which have common mechanisms with approved drugs for these diseases were identified by our analysis. Raloxifen (Selective estrogen receptor modulator), approved for the treatment of osteoporosis and Bazedoxifene (a newer generation of selective estrogen receptor modulators), approved for prevention of postmenopausal osteoporosis were both found to have opposite signatures to that of Cluster-PD. Other examples include metixene, an anticholinergic drug - already approved as antiparkinsonian, as well as other anticholinergic drugs, such as: solifenacin (approved for urinary incontinence), mebeverine (approved as antispastic) and deltaline. Metergoline, a dopamine receptor agonist and Darapladib (phospholipase inhibitor), an investigational drug currently in a clinical study for stabilization of atherosclerotic plaque, were also revealed in the analysis.

## Concluding remarks

The meta-analysis of human microarray datasets relevant to healthy and pathological aging led to a clear clustering of the transcriptional transitions between such biological states. For the available datasets, encompassing aging and age-related diseases, two main clusters of transcriptional shifts were identified - one corresponding mainly to changes reported in Alzheimer’s Disease and one for Parkinson’s Disease. Interestingly, amongst the drugs that could trigger a reverse transition to the consensus signatures of each cluster we identified several with strong links to the aging process, suggesting that the model and used methodology could have the potential to identify valuable candidate therapies and drugs for various pathologies. Although our application focused only on two clusters (in this case, both relevant for neurodegenerative diseases), this method can be extended to any pathology associated with aging. The impact of this approach should also increase with the accumulation of a larger and more diverse volume of data.

## Materials and methods

### Dataset selection and curation

The search for datasets was performed in two stages: 1) a programmatic search, using GEOmetadb, and querying it for relevant keywords (such as aging and aging, longevity, lifespan, etc), and 2) a manual curation stage, in which the description of the datasets, as well as in some cases the abstracts/body of the associated papers were carefully screened by our group. Dataset samples from nonagenarians, centenarians and supercentenarians, as well as from children or individuals in early stages of development (ex: fetal samples, testing induced pluripotency) were excluded from the study. For pathology-related datasets, samples from patients who were not under normal conditions (including those under treatment, diets, etc) were also excluded.

Additionally, for reducing noise generated by the variety of array platforms, normalizations and methodologies, the datasets with the most uncommon platforms were also discarded before the processing phase (in the end only microarray platforms that can be processed with R/Bioconductor scripts were kept). Datasets lacking relevant information (subject age unspecified, too broad age interval, etc), or considered inadequate for the study (due to insufficient sample size, lack of control, insufficient link to aging) were also discarded.

Age or age group for the samples was inferred using the *characteristics_ch1* field in GEOmetadb, and filled in manually where this value was missing. Tissue meta-data was also extracted from the data sample if possible, or filled in manually if required.

### Gene expression processing

Bioinformatics processing of all gene expression differences was performed from scratch, in a uniform manner for all datasets. Microarrays from the GEO database were retrieved, and analyzed using in-house R/Bioconductor-based scripts. Affymetrix datasets were normalized using the RMA method from the *affy* Bioconductor package and datasets which included other platforms were normalized using the quantile method from the *limma* package. Differential expression was computed for all comparisons of interest, given that there are at least 2 samples in each of the compared states. Benjamini-Hochberg multiple testing correction was applied for each comparison. If not otherwise specified in the text (such as for the custom meta-analysis, see below), genes were considered differentially expressed for an adjusted p-value < 0.05 and a |logFC|>1.

### The similarity score between 2 dataset comparisons

To calculate the similarity of the transcriptional changes that can be observed in comparisons from the same or different datasets, we have developed a Python script that computes a similarity score, based on the algorithm developed by Connectivity Map (Lamb et al., 2006; https://clue.io/connectopedia/pdf/cmap_algorithms). Briefly, the script performs a Gene Set Enrichment Analysis (GSEA) for a genetic signature (ex: a list of significantly changed transcripts), by quantifying similar modifications in terms of up- and down-regulation. Based on GSEA, two enrichment scores were computed (ES_up_ and ES_down_) and then the Weighted Connectivity Score (WTCS) was computed as (ES_up_ - ES_down_)/2 if sign(ES_up_) ≠ sign(ES_down_), 0 otherwise. The graph presented in Fig. 1, is constructed using edges as similarity scores.

#### Network analysis and visualization

The construction of networks and their visualization was performed using Cytoscape ver. 3.8.2. Clustering was performed with the MCL algorithm, using granularity = 5.

#### Meta-analysis of cluster data and Connectivity Map

For the meta-analysis, all expression change values corresponding to a p-value < 0.05 were taken from individual dataset DE analyses, without considering FDR. For each gene, N_up was computed as the number of dataset comparisons in which p-value < 0.05 and logFC > 0, while N_down was computed as the number of dataset comparisons in which p-value < 0.05 and logFC < 0 (the consistency criterion). In the cluster consensus signature, a gene was then considered upregulated if N_up > 0 and N_down = 0, and downregulated if N_down > 0 and N_up = 0. For the output of the meta-analysis, only genes with |logFC| > 1 (magnitude of the effect being at least 2x) were considered.

For the ConnectivityMap analysis, the 150 topmost up- and down-regulated genes from the meta-analysis were used as input. These were input in the CMap webtool, and the results were then manually analyzed. Sorting of the input genes, was done descending by max (|logFC|) across all studies in the cluster.

#### Functional module analysis

The construction of a network with functional modules for clusters of differentially-expressed genes was performed using the HumanBase tool (Greene et al., 2015), https://hb.flatironinstitute.org, with a minimum module size set to at least 10 genes. Briefly, HumanBase provides the possibility to identify, at the tissue level, functional modules containing genes and their interaction partners which specifically work together, by grouping them into clusters of relevant biological processes. HumanBase detects modules of genes from tissue-specific functional association gene networks built by integrating vast omics datasets and associates terms (e.g. processes, pathways) to the detected modules based on overrepresentation. In the functional modules analysis, all networks were based on the global network while using brain tissue or nervous system tissue-based networks the results were extremely similar.

#### Enrichment analysis

The GSEA – prerank analysis was performed using python, with the gseapy library wrapper of GSEA (Mootha et al., 2003; Subramanian et al., 2005). The analysis is easily reproducible using the data and code from https://github.com/ursueugen/transcriptional-profiles_enrichment-analysis (public repository). Briefly, we run prerank analysis on the data obtained from the meta-analyses with 1000 number of permutations with respect to common functional gene sets collections that we pre-selected: ‘KEGG_2021_Human’, ‘GO_Biological_Process_2021’, ‘Reactome_2022’, ‘ChEA_2022’. For post-processing and summarizing the results, we examined the significant terms (FDR-adjusted p-value <0.01) with the highest absolute value of the normalized enrichment score (NES).

## Acknowledgments

This work was supported by the Romanian Ministry of Education and Research, CCCDI - UEFISCDI, through PNCDI III [Grant numbers: PN-III-P1-1.1-TE-2019-1020 and PN-III-P2-2.1-PED-2019-2593 to RT]. We are also grateful for the funding received from the Dr. Amir Abramovich Research Fund [granted to VEF].

## Author contributions

This study was carried out by RT’s research groups, in collaboration with VEF. Data collection, processing, analysis of the result and their description were done by GB, DT and EU. Interpretation of the results was done by all authors. RT designed, coordinated and supervised the project. VEF and SG critically reviewed the project and the results. All authors have participated in the writing of the manuscript. All authors reviewed the manuscript.

